# A Head and Neck Cancer Patient-Specific Microphysiological System for Predicting Response to Chemoradiation

**DOI:** 10.64898/2026.04.28.721391

**Authors:** Adeel Ahmed, Nathan Hendrikse, Rene Welch Schwartz, Yuanshan Li, Marcos Lares, Cassidy K. Felix, Adam R. Burr, Irene M. Ong, Paul M. Harari, David J. Beebe, Sheena C. Kerr

## Abstract

Head and neck cancer (HNC) is the 6th most common malignancy worldwide. 60% of patients present with advanced disease and approximately 50% of patients recur following primary treatment. Chemoradiation remains a standard of care for most patients. However, clinicians lack functional tools to predict which patients will respond to chemoradiation prior to treatment and current models, including organoids and animal model systems, fail to capture either full complexity or patient-to-patient heterogeneity of the individual HNC tumor and microenvironment (TME). Here, we have developed, characterized, and tested a patient-specific microphysiological system (MPS) that reconstructs the HNC TME in a vascularized 3D environment. This MPS was constructed from malignant cells, fibroblasts, and immune cells from a patient’s surgically resected tumor, seeded within a 3D hydrogel with molded endothelial lumens. Single-cell RNA sequencing confirmed that the MPS preserved 12 transcriptionally distinct cell populations found in matched native tissue. The platform recapitulated tumor hypoxia, with a 12-fold increase in hypoxic marker expression that altered radiation response, consistent with clinical HNC biology. Compartment-resolved imaging revealed distinct treatment dynamics in tumor, stromal, and vascular regions, and individual patients exhibited divergent responses to chemoradiation in spheroid morphology, cell viability, and migration. We found the slope of spheroid area change with treatment tracked with tumor recurrence, suggesting this metric could serve as a functional predictor of therapeutic response.

## Introduction

Head and neck cancer remains the 6th most common malignancy globally and poses unique clinical challenges pertaining to patient stratification for therapeutic selection^1^. 60% of patients present with advanced disease, and half of these patients will experience recurrence^2,3^. The current standard of care includes some combination of surgery, radiation, chemotherapy or neoadjuvant immunotherapy depending on patient risk group. Given the advanced nature of the disease and the choice of multiple treatment modalities with significant associated risks, any mechanism to stratify patients into treatment groups based on resistance and response profiles will improve both patient outcomes and reduce toxicity, resulting in an overall improvement in the quality of care^4–7^.

The HNC tumor microenvironment (TME) modulates tumor progression, invasion, and response to treatment^8–10^ through a coordinated network of biophysical and biochemical interactions between malignant cells, fibroblasts, vasculature, immune populations, and the physical extracellular matrix (ECM)^11–14^. Current tumor profiling methods - both molecular and functional - such as RNA sequencing or in vitro organoids cannot fully dissect the complex interactions in the TME that are driving malignant processes in the tumor. Moreover, recent evidence has highlighted significant patient-to-patient heterogeneity, which cannot be fully recapitulated by animal or current in vitro models, limiting our ability to investigate drug response profiles at the individual level in a spatiotemporal manner across the TME. Thus, it follows that an ideal platform to evaluate tumor response to therapy should be able to capture the complexity of the human TME, enable easy perturbation and investigation, respond to and predict clinical treatment outcomes, all while also being amenable for laboratory execution for individual cases.

Microphysiological systems (MPS) are a class of microfluidic in vitro culture platforms that exploit microscale phenomena to recapitulate the biological and physical complexity of tissue at the cellular scale to recreate in vivo-like tissue environments, in vitro^15–17^. We have already shown the development of the LumeNEXT platform^18^ - a 3D cell culture environment with vascularized lumens - to investigate tumor cell migration^10,19–21^, personalized response of breast cancer to therapy^22,23^, and study interactions between different components of the TME. Our platform is able to consistently recreate a 3D luminal construct with defined size and geometry within a 3D collagen hydrogel embedded with stromal TME components and captures endothelial biology with higher fidelity than 2D or 3D cultures by enhancing cytokine secretion, endothelial barrier formation, and providing a full 3D matrix^24^. Moreover, we have also shown that the HNC TME is sensitive to the presence of lymphatic vessels which can promote tumor cell migration^19^.

Based on these data, we posit that a LumeNEXTLumeNEXT MPS which incorporates HNC cells from a single patient’s tumor, within 3D collagen hydrogels with both vascular and lymphatic lumens, will provide a viable model for evaluating patient-specific treatment responses across the TME using functional readouts. The LumeNEXT MPS enables co-culture of five matched cell populations extracted from surgically resected HNC tissue using a digest protocol that preserves the tumor infiltrating leukocyte (TIL) population and is co-cultured in an optimal media formulation. Using single cell RNA sequencing (scRNA-seq), we demonstrate that the cell populations in the model exhibit significant overlap with those in the native tissue. We also show our HNC model captures hypoxia, a hallmark of HNC^25–27^, and responds to radiation and chemotherapy. Additionally, the model can recapitulate patient-specific differences in cell migration, angiogenesis, and cellular morphology. Lastly, we show that this patient-specific MPS captures patient heterogeneity in response to chemoradiation, which can be correlated with patient clinical data. Together, these data highlight the potential of MPS as a powerful tool for patient stratification and precision functional oncology workflows.

## Materials and Methods

### Tissue Acquisition

Head and neck cancer tissue was obtained from surgical resections upon informed consent from patients (IRB protocols 2021-0312 and 2023-1237) at the UW Health Hospital, University of Wisconsin, Madison, WI. After resection, tissue was immediately placed in a specimen container with 20mL transport media containing DMEM + 4.5g L^−1^ D-glucose (Thermo, 11965092), 1% (v/v) penicillin/streptomycin (Gibco, 15140122), 10% (v/v) fetal bovine serum (Avantor, 76419584), 400ng mL^−1^ hydrocortisone (Sigma, H6909), 5 µg mL^−1^ insulin solution from bovine pancreas (Sigma, I0516), 1%(v/v) nystatin (Sigma, N9150), and 1%(v/v) amphotericin B (Sigma, A2942). Tissue was stored in the transport medium at 4 °C for no more than 24 hours after collection.

### Tissue Digestion

Tissue was weighed and a 4 mm biopsy was collected for scRNA-seq and flash frozen in liquid nitrogen. The remaining tissue was placed in a 100 mm petri-dish (non-cell culture treated) on ice. 3 mL of digest media was added (transport media, 1X collagenase/hyaluronidase (STEMCELL Technologies, 07912)) to the dish. The tissue was then cut into small pieces using scalpels until each piece was less than 1 mm^3^ in size. The tissue pieces were equally distributed into two 15 mL centrifuge tubes into which 3 mL of digest media was added. Into one of the tubes, 0.1%(v/v) dispase (Worthington Biochemical Corporation, LS02104) was added to enhance the digestion and release of epithelial cells however the viability of the immune population is greatly reduced. The remaining tube containing 3 mL 1X collagenase/hyaluronidase was designated for the isolation of tumor infiltrating leukocytes (TILs).

The tubes were placed in a rotisserie hybridization incubator at 37°C for 2 hours. The digested tissue was strained through a 70 µm strainer to remove undigested pieces of tissue. The cells were collected, counted and washed using either Pneumacult EX-Plus (StemCell Technologies, 05040) for epithelial culture or Fibroblast Media (ScienCell, 2301) for fibroblast culture before being split into T12.5 flasks for propagation and freezing. TILs were isolated from tissue using REALease CD45 (TIL) MicroBeads, human kit (Miltenyi Biotec 130-121-563) and cryopreserved in cryopreservation media (Sigma-Aldrich, C2874-100ML).

### Co-Culture for Media Screening

MicroDuo well plates (Onexio Biosystems) were used to co-culture different cell types for media screening. Within a pair of adjacent wells, the wells are separeated by a short barrier. When each well is filled with 50µL of media, adjacent wells are disconnected and maintain a monoculture, and when media volume is increased to 100µL per well, the media bridges across the short barrier to form a connected co-culture. To evaluate cell viability under different media formulations and co-cultures, cells were seeded at a density of 10,000 cells cm^2^ and monocultures in default cell-type specific media were established for 24 hours. Cells were bridged to establish a co-culture after 24 hours, which was maintained for 72 hours. Cell density and coverage were optically verified under a light microscope and viability was measured using a luminescence assay (Promega CellTiter-Glo G7570). Briefly, CellTiter-Glo was mixed 1:1 with serum-free basal media (RPMI 1640, Gibco, 11875093) and added to cell culture after aspirating culture media. Cells were shaken on an orbital shaker for 5 minutes at 500 RPM, followed by a 10-minute incubation at room temperature. Luminescence was read out no more than 30 minutes after adding the CellTiter-Glo using a plate reader (Pherastar FS, BMG Labtech).

### Cell Culture and Expansion

UM-SCC1 cells were cultured with DMEM high glucose (Gibco, 11964-092) and supplemented with Penicillin/Streptomycin (Gibco, 15140), 10% FBS (VWR, Randor), and MEM Non-Essential Amino Acids (Gibco, 11140-050). Human umbilical vein endothelial cells (Lonza C2519A) were cultured with endothelial basal medium-2 (Lonza, CC-3156) and supplemented with EGM-2 SingleQuot Kit (Lonza, CC-3162) and Penicillin/Streptomycin. Human lymphatic endothelial cells (Sciencell, 2500) were cultured with endothelial cell media (ScienceCell, 1001) and supplemented with HLEC comprehensive kit (ScienceCell, 2500). Patient-derived epithelial cells were cultured with PneumaCult-Ex Plus Media (Stemcell, 05041) and supplemented with PneumaCult-Ex Plus 50X supplement (Stemcell, 05042) which helps retain HNC epithelial cell characteristics^28^. Patient-derived fibroblasts were cultured with Fibroblast Media complete kit (ScienceCell, 2301). Patient tumor infiltrating lymphocytes (TILs) were cultured with RPMI Media (Gibco, 11875135) and supplemented with HEPES,

Human AB Serum, L-Glutamine, Penicillin/Streptomycin, and IL-2. All cells were cultured in the MPS using a combination of fibroblast media, endothelial basal medium-2, and Pneumacult media, and their respective supplements. Cells were grown at 37 °C with 5% CO2, and the media was refreshed every 2-3 days.

### 3D cell culture

To create maximum hydrogel adhesion to the device, 2% poly(ethyleneimine) (PEI, Sigma-Aldrich, 03880) was diluted in deionized water, loaded into the chamber through side ports, and incubated at room temperature for 15 minutes. The PEI was then aspirated and replaced with 0.4% glutaraldehyde (GA, Sigma-Aldrich, G6257) also diluted in deionized water and incubated at room temperature for 45 minutes. Following incubation, devices were washed three times with sterile deionized water to remove residual GA and were subsequently ready for ECM loading.

The ECM solution was prepared by diluting high-density rat-tail collagen type 1 (Corning, 354249) with 10X PBS, fibronectin bovine plasma (Millipore Sigma, F1141), and media, and then neutralized with 1M NaOH (Fisher Scientific, S318) to achieve a final concentration of 4 mg mL^−1.^ For experiments that included fibroblasts and TILs, a final concentration of 1000 cells µL^−1^ was added to the media dilution to maintain the same final overall concentration. Spheroids were then collected and gently mixed into the ECM. A 13 µL volume of ECM containing 3-4 spheroids was injected into the device side port and incubated at room temperature for 30 minutes. To minimize evaporation, 200µL of DI water was placed in the surrounding dish. During incubation, both HUVECs and HLECs were passaged using 0.05% trypsin-EDTA (Gibco, 25300-062), neutralized with complete media, centrifuged, and resuspended at 20,000 cells µL^−1^. PDMS rods were carefully pulled out of the small port using a sterilized tweezer. After creating two hollow lumens, HUVECs (2 µL) were added to the left lumen and HLECs (2 µL) were added to the right lumen at 20,000 cells µL^−1^. After seeding, devices were incubated for 1 hour at 37 °C with manual flipping every 15 minutes to ensure uniform cell attachment throughout the lumen. Finally, 10 µL of head and neck (HNC) media was added to each lumen, and lumens were cultured overnight at 37 °C. Media was changed daily using HNC media supplemented with IL-2 (1 µL mL^−1^).

### Spheroid Generation

Tumor spheroids were generated via the hanging drop method as previously described_19_. In short, UM-SCC-1 or patient derived epithelial cells were trypsinized, counted, and resuspended at 6,000 cells µL^−1^ in media containing methylcellulose. Droplets (25 µL) were then displaced onto the lid of a cell culture plate (Thermofisher, 242811) with 20mL of distilled water added to the bottom of the dish to minimize evaporation. After 24 hours of incubation at 37 °C, a single spheroid formed in each droplet which were used within 3 days.

### Cisplatin and Radiation Treatment

Cisplatin (SelleckChem, S1166, USA) was diluted in culture media and delivered into the MPS via the vascular lumens. Post treatment, Cisplatin was washed out with neat culture medium with 3 washes for 15 minutes each.

Radiation was carried out using a cell-irradiator (Xstrahl, USA) at a distance of 30mm, at 195kV at fractions and doses mentioned in each experiment. Each fraction was spaced 24 hours apart.

### Hypoxia visualization

Hypoxia was generated at 1% O_2_ within a cell culture incubator using a gas flow controller (Coy Lab Products, MI, USA). For the 2D condition, cells were plated at a density of 10,000 cells cm^−2^ in a 96 well plate and transferred to the incubator via an airlock to ensure hypoxia was maintained. Spheroids and MPS were developed as described above. Cells were incubated in the chamber for 24 hours with Image-iT Hypoxia Green reagent (Invitrogen, I14833) at 5µM concentration. Cell culture dishes were sealed with parafilm immediately upon removal from the chamber to minimize O_2_ equilibration and imaged within 1 hour of removal from the chamber.

### Imaging

Prior to spheroid formation, UM-SCC-1/patient-derived epithelial cells were stained with a long-lasting cell membrane dye (Vybrant DiD, ThermoFisher, V22887) at 1:40 in media for 10 minutes. Cells maintained staining for the duration of the experiments (9 days). For measurement of cell viability, cells were stained with Calcein-AM (Invitrogen C1430), propidium iodide (Molecular Probes, P1304MP) and Hoechst 33342 (Thermo Scientific, 62249) in serum free media for 30 minutes, followed by three 15-minute washes. Confocal Z-stack imaging was performed using a Nikon Ti-2 Eclipse confocal with a Yokogawa Spinning Disk and processed using an automated segmentation and quantification pipeline implemented in Python using scikit-image, StarDist^29–31^, and OpenCV.

### Image Processing

Confocal Z-stacks were processed using an automated image processing pipeline implemented in Python 3.11 (Python Software Foundation) with scikit-image for morphological analysis. Each Z-stack was max-projected and segmented into three compartments: lumens, spheroids, and stroma. Individual nuclei were identified using the StarDist deep learning-based nuclei segmentation model^29–31^. Tumor cells were distinguished from stromal populations by DiD membrane dye fluorescence. Quality control filters were applied to exclude imaging artifacts and debris: spheroids with segmented areas below 10,000 pixels were excluded from analysis, and data from the patient sample were filtered separately from SCC1 cell line controls to ensure patient-specific analyses. The pipeline extracted 76 morphological and intensity-based features per spheroid, including shape descriptors (area, perimeter, eccentricity, solidity, major/minor axis lengths), texture features, and multi-channel fluorescence intensities (DAPI, propidium iodide, Calcein AM). For this study, four primary readouts were analyzed: spheroid area, cell viability, nuclei count, and nuclei size. Cell viability was quantified as the ratio of propidium iodide-positive nuclei to total DAPI-positive nuclei within each compartment, expressed as a percentage. Nuclei area was used as an indicator of cell health, with decreased area associated with apoptotic morphology. Spheroid area was measured from the segmented spheroid boundaries in each projected image.

### Tissue Fixation and Preservation for scRNA-seq

Tissue fixation and dissociation were performed according to 10x Genomics Fixed RNA protocols. For three patient samples, tissue fixation and dissociation were performed according to the Chromium Next GEM Single Cell Fixed RNA Sample Preparation protocol (CG000553) using the Chromium Next GEM Single Cell Fixed RNA Sample Preparation Kit (10x Genomics, PN-1000414). For one additional patient sample, tissue fixation and dissociation were performed using the newer GEM-X Flex sample preparation protocol (CG000783) with the GEM-X Flex Sample Preparation v2 Kit (10x Genomics, PN-1000781). Briefly, flash-frozen patient tumor tissue was minced on a pre-chilled glass dish on ice and fixed in Fixation Buffer B for 16–24h at 4 °C. Following fixation, samples were washed with chilled PBS and resuspended in Quenching Buffer B. Fixed tissue pellets were dissociated in pre-warmed RPMI containing Liberase TH (1 mg/mL; MilliporeSigma, Cat# 5401135001) and incubated at 37 °C for 40 min using an Octo Dissociator (Miltenyi Biotec). Dissociated samples were passed through a 30 µm filter and stored at −80 °C until further processing.

To recover cells from MPS devices, samples were incubated in collagenase P (8 mg/mL; MilliporeSigma, Cat# 11213857001) for 10 min at room temperature to digest the collagen matrix. Samples were centrifuged at 300 × g for 10 min, resuspended in 0.05% trypsin, and incubated at 37 °C for 5 min to generate a single-cell suspension. After centrifugation, cell pellets were fixed using the Chromium Next GEM Single Cell Fixed RNA Sample Preparation Kit or GEM-X Flex Sample Preparation v2 Kit and stored at −80 °C.

### scRNA-seq Library Preparation

Library preparation was performed according to the appropriate 10x Genomics Fixed RNA profiling workflow for each chemistry. Three patient samples were processed using the Chromium Next GEM Single Cell Fixed RNA Hybridization & Library Kit (PN-1000415), Chromium Next GEM Single Cell Fixed RNA Human Transcriptome Probe Kit (PN-1000420), Chromium Next GEM Single Cell Fixed RNA Gel Bead Kit (PN-1000421), Chromium Next GEM Chip Q Single Cell Kit (PN-1000422), and Dual Index Kit TS Set A (PN-1000251). One additional patient sample was processed using the GEM-X Flex v2 workflow with GEM-X Flex GEM & Library Kit (PN-1000782), GEM-X Flex Hybridization & Wash Kit (PN-1000789), GEM-X Flex Human Transcriptome Probe Kit (PN-1000785), GEM-X Flex Gel Bead Kit (PN-1000790), GEM-X Flex Gene Expression Chip Kit (PN-1000791) and Dual Index Kit TS Set A (PN-1000251). Fixed single-cell suspensions were loaded onto a Chromium iX instrument to generate gel beads-in-emulsion (GEMs) and construct barcoded cDNA libraries. Libraries were assessed using High Sensitivity D5000 ScreenTape (Agilent, PN-5067-5592) with High Sensitivity D5000 Reagents (Agilent, 5067-5593) on a TapeStation and sequenced on a NovaSeq X Plus (Illumina) using paired-end, dual-index reads (28 bp Read 1, 10 bp i7, 10 bp i5, 90 bp Read 2) with a 10% PhiX spike-in, following 10x Genomics recommendations. Sequencing was performed to a target depth of 50,000 reads per cell.

### Single-cell RNA-sequencing

Single cell RNA-seq was performed. Cellranger (version 9.0.1) with the GRCh38-2024A reference and the Chromium Human Transcriptome probeset was used to demultiplex, barcode processing and quantify gene expression. We filtered low quality cells based on the following quality metrics: total # of UMIs per cell, # of detected genes, defined as the number of genes with positive expression per cell, and mitochondrial content. We filtered out cells with less than 1K total UMIs, with less than 300 detected genes and more than 20% mitochondrial content. We kept tissue cells with less than 60K UMIs and MPS cells with less than 250 UMIs. Expression was normalized using scran (version 1.38)^32^ and then converted to log normalized expression values. Tissue and MPS samples were integrated separately using Harmony (version 1.24.0)^33^ with at most 20 clusters.

Tissue and MPS integrated datasets were analyzed using Seurat (version 5.4.0)^34^ and used the Louvain clustering algorithm^35^ with resolution=1, and then were annotated by examining the expression of the gene markers in **Table 1**. Tissue and MPS single cell integrated datasets were integrated using STACAS (version 2.4.1)^36^ and these annotations as cell labels for semi-supervised integration with parameters: k.anchor=50, k.score=30, anchor.coverage=30% and label confidence=75%. Uniform Manifold Approximation and Projection (UMAP)^37^ representations were calculated using the top 25 principal components and 100 neighbors. The UCell R package (version 2.14)^38^ was used to compute U-statistic based module scores for the cell type gene sets in **Table 1**; and these scores were utilized to annotate the integrated datasets clusters, obtained with the Louvain algorithm with resolution=1.

**Table 1.**
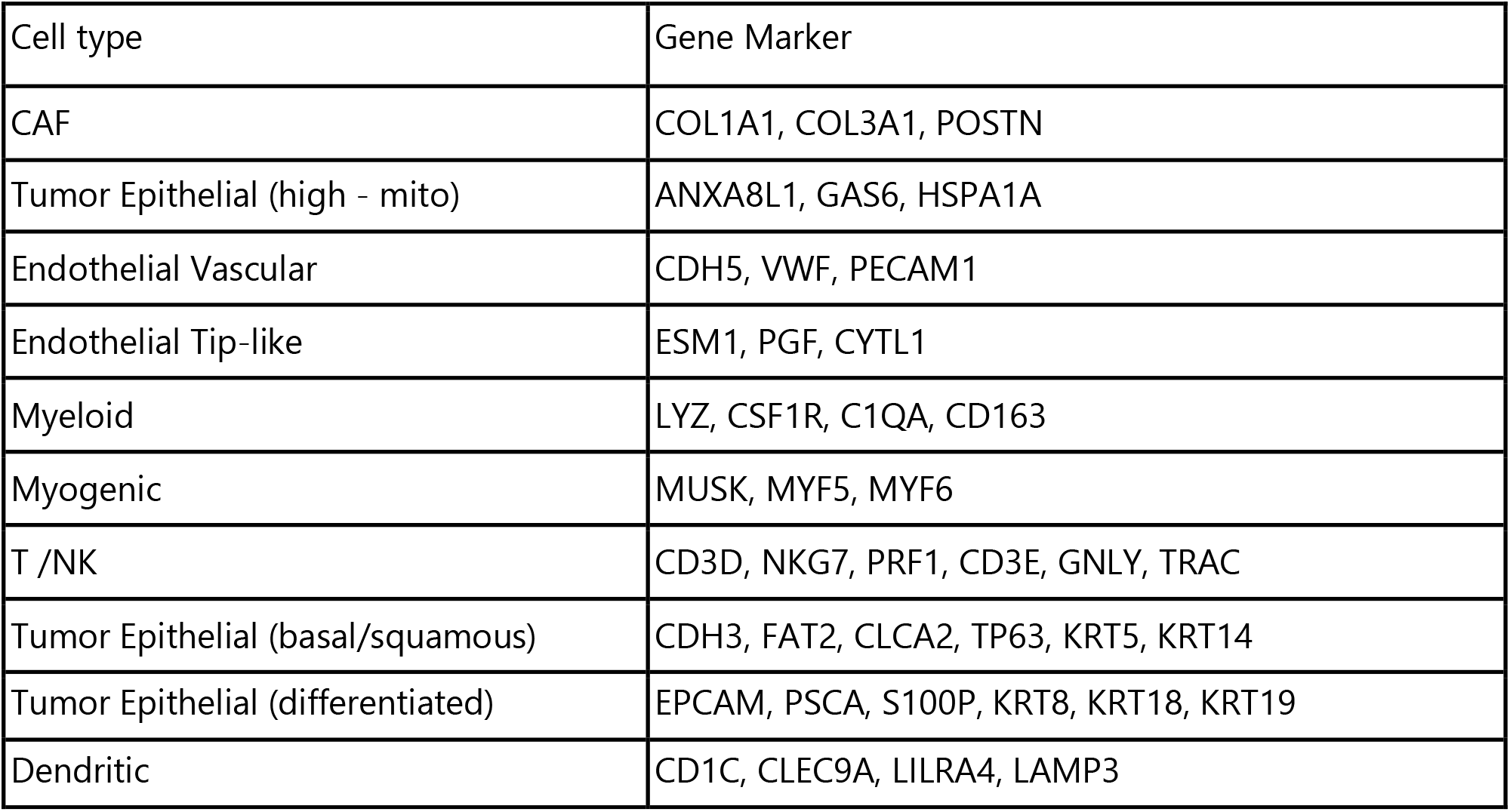
Table listing markers used for cell type annotation.

### Statistical Analysis

Statistical analyses were performed in R, using ggplot2 and ggpubr packages for data visualization. Data were normalized using two approaches: (1) fold-change relative to untreated controls (Gy=0, CSP=0) for each metric, and (2) Z-score normalization (Glass’s Delta: (metric - control_mean) / control_std) to standardize responses across patients. Kruskal-Wallis test was used to test for radiation dose effects; otherwise, Post-hoc pairwise comparisons were performed using Tukey’s HSD test (parametric) or Dunn’s test with Bonferroni correction (non-parametric). Effect sizes were calculated using Cohen’s d for pairwise comparisons and eta-squared (η^2^) for omnibus tests. Two-way ANOVA was used to test for interaction effects between radiation dose (Gy) and cisplatin concentration (CSP) on treatment response metrics, with partial eta-squared reported as the effect size. Dose-response relationships were characterized by fitting linear regression models to spheroid area, cell viability, and cell count as a function of treatment intensity for each patient and cisplatin condition. The resulting slopes were used as summary metrics of treatment sensitivity. Data are presented as mean ± standard deviation unless otherwise noted, and a significance threshold of p < 0.05 was used throughout.

## Results

### Construction of Head and Neck Cancer Microphysiological System

The HNC TME comprises heterogeneous populations of malignant epithelial cells, cancer-associated fibroblasts (CAFs), blood endothelium, lymphatic vessels, and immune cells. In vivo, these cells are surrounded by a collagen-rich ECM that provides both structural and biochemical support to the tumor. To generate the HNC TME, we integrated cells obtained from surgically resected tumor specimens into the LumeNEXT MPS. First, the tissue was obtained from routine surgical resection of the tumor prior to treatment. As shown in **Figure 1A**, the tumor tissue was split into two pieces. The first piece was minced and digested using collagenase, hyaluronidase and dispase to release fibroblasts and epithelial cells within the tissue, but with reduced TIL viability. After tumor tissue was digested, the cell suspension was filtered to remove debris, and the cleared suspension was seeded into fibroblast- or epithelial-specific medium to selectively grow both cell populations. The second tissue piece was digested without dispase but with collagenase and hyaluronidase and TILs were isolated using a CD45+ bead-based isolation and immediately frozen.

**Figure 1.**
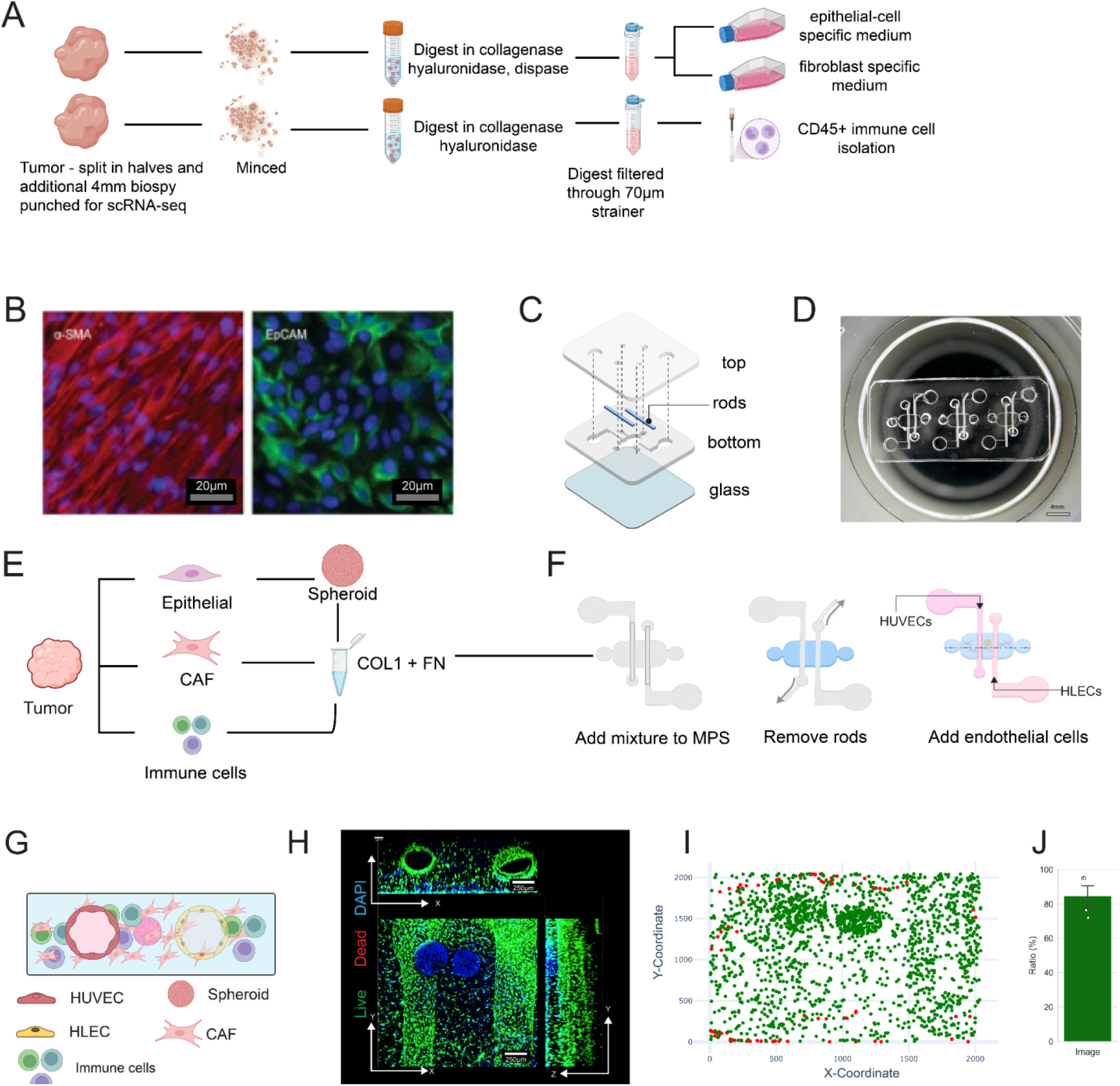
(A) Schematic showing the workflow for isolating cells from a single piece of tumor tissue. (B) alpha-SMA and EpCAM staining to validate cancer associated fibroblast (CAF) and epithelial phenotype. (C) Schematic showing the assembly of the LumeNext dual lumen MPS platform. (D) Actual image of the LumeNext dual lumen MPS. (E-F) Schematic depiction of the workflow to load patient-tumor cells into a 3D collagen hydrogel and the MPS. (G) Schematic cross-section view of the MPS. (H) Confocal micrograph showing cell viability in the MPS after 5 days of culture. (I-J) Dot plot representing the spatial distribution of live and dead cells in the MPS and cell viability across 3 devices after 5 days of culture.

Following expansion, alpha-SMA and EpCAM were used as canonical fibroblast and epithelial markers to validate the cell phenotype in addition to morphological assessments **(Figure 1B)**. We have previously validated the phenotypes of fibroblasts extracted using this protocol via qPCR for canonical markers ACTA2, FAP, Vimentin, COL1A1 and FSP1^19^. The epithelial cells were formed into three-dimensional spheroids containing 2000 cells each. Before loading into the MPS, the tumor spheroids, CAFs, and immune cells were mixed into a 4 mg mL^−1^ collagen fibronectin hydrogel.

To recreate the HNC TME in an MPS in vitro, we developed an MPS based on the previously published LumeNEXT platform^18^. The LumeNEXT uses micromolding to create luminal structures within a cell-laden, collagen-fibronectin hydrogel. This platform consists of two PDMS sheets bonded together to form a cell culture chamber (2 mm x 3 mm x 0.75 mm) **(Figure 1C-D)**. Within this chamber, two removable rods (280 µm) were placed as shown in **Figure 1D**. This construct was fixed on to a glass-bottom dish, enabling confocal imaging of the engineered HNC TME.

The cellular components were loaded sequentially **(Figure 1E)**: initially, collagen-embedded epithelial spheroids, CAFs, and immune cells were added to the culture chamber (Step 1). This collagen-cell mixture polymerized around the rods within the cell culture chamber, forming a 3D cell culture environment within the MPS. Following collagen polymerization, the PDMS rods within the chamber were removed using sterile tweezers. The rods left behind luminal structures within the collagen which could be perfused (Step 2). Subsequently, 40,000 human umbilical vein endothelial cells (HUVECs) and human lymphatic endothelial cells (HLECs) were introduced into the two lumens separately to form a physiologically relevant vascular environment. The small size of the patient’s tumors and the limited cell numbers precluded the isolation of endothelial or lymphatic cells from the patient tissue. The seeding density of endothelial cells was empirically determined by confocal microscopy to ensure maximal cell coverage within the lumens. (Figure S1). **Figure 1G** shows a cross-sectional view of the cell culture chamber in the MPS and the resulting cellular composition after the complete model had been established.

The successful spatial organization of the lumens and stromal cells was verified using confocal microscopy. Calcein AM and propidium iodide staining were used to assess the viability of cellular components 5 days post-culture within the MPS **(Figure 1H)**. Viability staining revealed robust cell viability throughout the construct, with DAPI-positive nuclei indicating cellular integrity and the absence of necrotic areas. Quantitative image analysis confirmed approximately 85% cell viability across all cell populations throughout the construct, validating the platform’s capacity to sustain complex multicellular interactions under physiologically relevant conditions **(Figure 1, I-J)**.

Our approach to developing an HNC MPS with primary cells thus resulted in an MPS that faithfully recreates the composition and structural motifs seen in HNC tissue in vivo.

### Cell viability and TIL phenotype are maintained in co-culture using a custom culture media formulation in the MPS

Within the HNC MPS, we sought to integrate tumor epithelial cells, fibroblasts, and endothelial cells in a 3D environment. Each of these cell populations demands specific nutrients, and the viability of primary cells is significantly affected by culture conditions. To maximize cell viability in co-culture, screening of several combinations of epithelial, fibroblast, and endothelial media with varying supplement fractions was carried out.

The microDUO plate (Onexio Biosystems) **(Figure 2A)** allows for monoculture in a pair of adjacent wells when the media volume is less than 50 µL per well. The short barrier between wells within the pair allows the cultures to be bridged with a media volume of 100 µL per well, allowing cells to be in co-culture by sharing soluble factors. This plate enabled the establishment of monocultures of fibroblasts, epithelial cells, or endothelial cells in individual wells of a pair, using native media for 24 hours, before bridging the cultures with custom media formulations to assess cell viability in co-culture.

**Figure 2.**
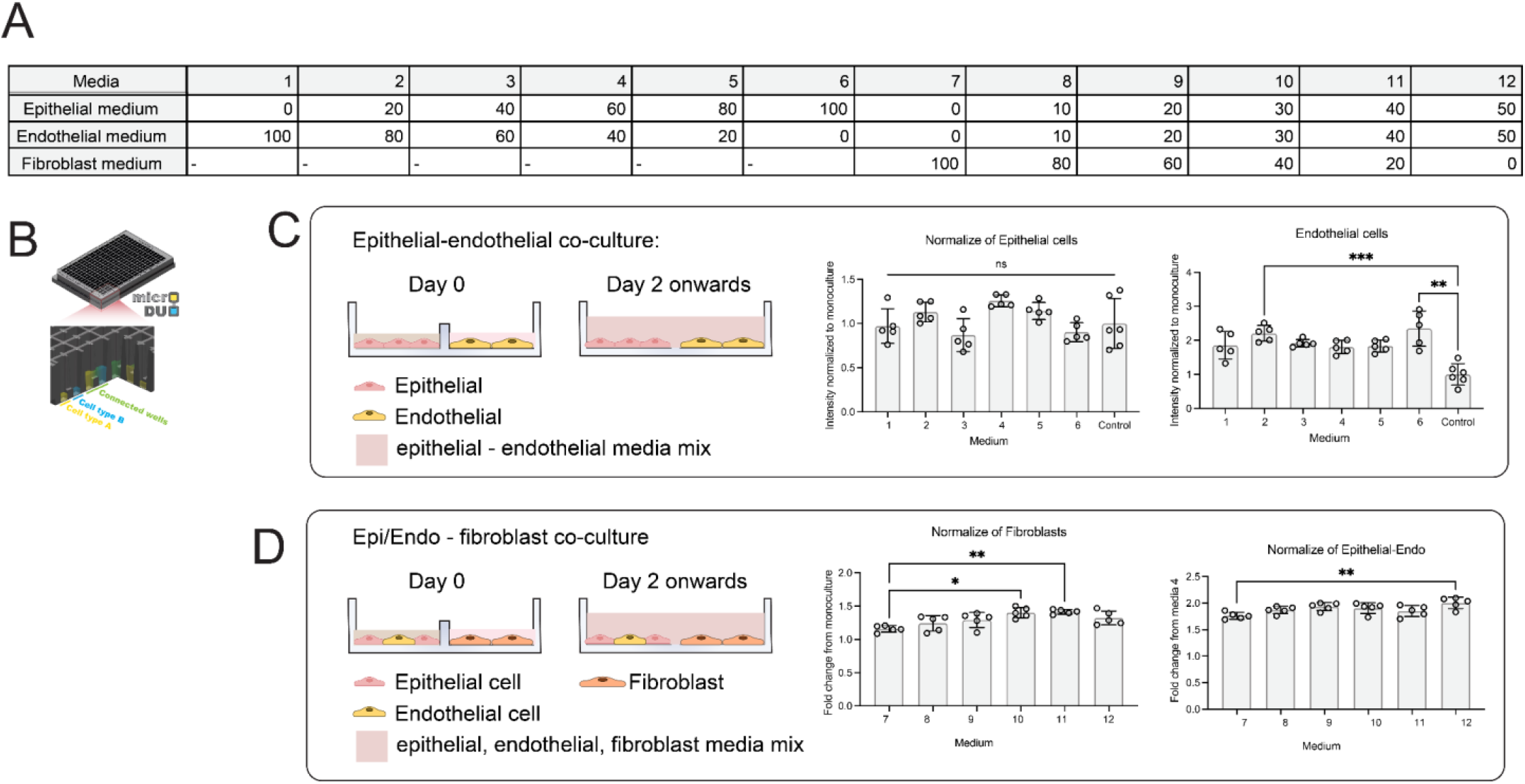
(A) Table showing the basal media ratios of different media formulations. All media formulations were fully supplemented with the provided manufacturer supplements. (B) Schematic cross-section of the microDUO plate showing wells separated at media volumes less than 50µL and connected at media volumes of 100µL per well. (C) Epithelial - endothelial co-culture viability across media formulations. (D) Epithelial/endothelial and fibroblast co-culture viability with different media formulations.

12 media formulations **(see Fig. S2 for full formulation)** were tested for maximum viability of epithelial cells, endothelial cells, and fibroblasts in co-culture. In the first set of experiments, cell viability was maximized for epithelial and endothelial co-culture because we observed that primary HNC epithelial cells are highly sensitive to their culture environment. As seen in **Figure 2B**, epithelial cell viability was not significantly affected by the media formulations. However, endothelial cell visibility was significantly higher in co-culture conditions with media 2 and media 6, showing over a two-fold increase in viability compared to native media control. Based on this data, we prioritized for epithelial cell viability, which was found to be highest in media 4 (1.26 fold increase over native media), and still provided increased viability for endothelial cells over control (1.92 fold increase over native media)

Next, the experiment was repeated with epithelial and endothelial cells co-cultured in a single well, and fibroblasts in an adjacent well. Fibroblasts were seeded in native media, whereas endothelial and epithelial cells were seeded in media 4. After 24 hours of culture, the media were bridged using 6 formulations comprising varying concentrations of fibroblast-, epithelial-, and endothelial-specific media supplements. Fibroblast viability was found to be highest in media 10 and 11 (1.40 and 1.42 fold increase over native media), whereas the epithelial-endothelial co-culture showed highest viability in media 4. However, the endothelial and epithelial compartment also showed increased viability across all fibroblast co-culture conditions, with fold-change increases ranging from 1.16 (media 7) to 1.42 (media 11). These data showed that while media formulations did have some significant impacts on cell proliferation, the effect size was generally small, and the co-culture environment was a stronger driver of cell proliferation. Therefore, we concluded that co-culture is not only beneficial for recreating the TME, but also improves cell viability for all populations. Additionally, based on these data, we selected media 10 (a formulation with equal parts of basal media with 100% supplements) for all the following experiments.

### scRNA-seq shows that HNC MPS maintains molecular similarity to the original tissues

The phenotypic trajectory of cells in vitro is strongly influenced by biophysical and biochemical cues including cell-cell interactions, cell-matrix interactions, and soluble factor concentrations. Given the environmental differences within a tumor tissue in vivo and our protocols for cell isolation, expansion and culture in vitro, there are multiple opportunities for phenotypic drift or losses in cell subpopulations. However, in a complex MPS with multiple cell-types, decoupling the effect of every environmental cue and the effect of these cues on cell phenotype is not practically feasible. Additionally, we are limited in our ability to profile the numerous variables which can potentially influence cell phenotype from tissue isolation, to expansion and subsequent co-culture in the MPS.

Therefore, to validate that the cells introduced within the HNC MPS continue to maintain phenotypic concordance with the matched patient tissue of origin, we employed scRNA-seq to compare the transcriptional similarity between the MPS and a biopsy of the matched primary tumor tissue from the patient. Surgically resected tissue was biopsied and flash frozen to preserve cells, while the rest of the tissue was digested to extract epithelial cells and fibroblasts. After expansion, MPS co-cultures were established and maintained for 24 hours prior to scRNA-seq using the 10X Genomics NEXT-GEM Flex kit.

The data from patient tissue and MPS were integrated using a two-step process: i) The subject samples for the tissue and the MPS datasets were integrated using Harmony to ensure that the subject diversity at each cluster is maximized and that way avoid annotating subject specific cell types; ii) These tissue and MPS integrated datasets were integrated with the semi-supervised version of STACAS using the annotated Harmony clusters as prior cell type information. STACAS is an extension of Seurat reciprocal PCA integration method tailored to heterogeneous single cell RNA-seq datasets that utilizes prior cell type annotations to guide the integration, and preserve the sample specific biological variation in the integrated dataset. UMAP visualization of the integrated datasets demonstrated a high degree of transcriptomic concordance between MPS-derived and patient-derived cells, with co-localization of cells from both sources **(Figure. 3A)**. This overlap was consistent across all patients, indicating that the MPS preserves the transcriptional landscape of the matched patient tissue and that this fidelity is not patient-specific.

**Figure 3.**
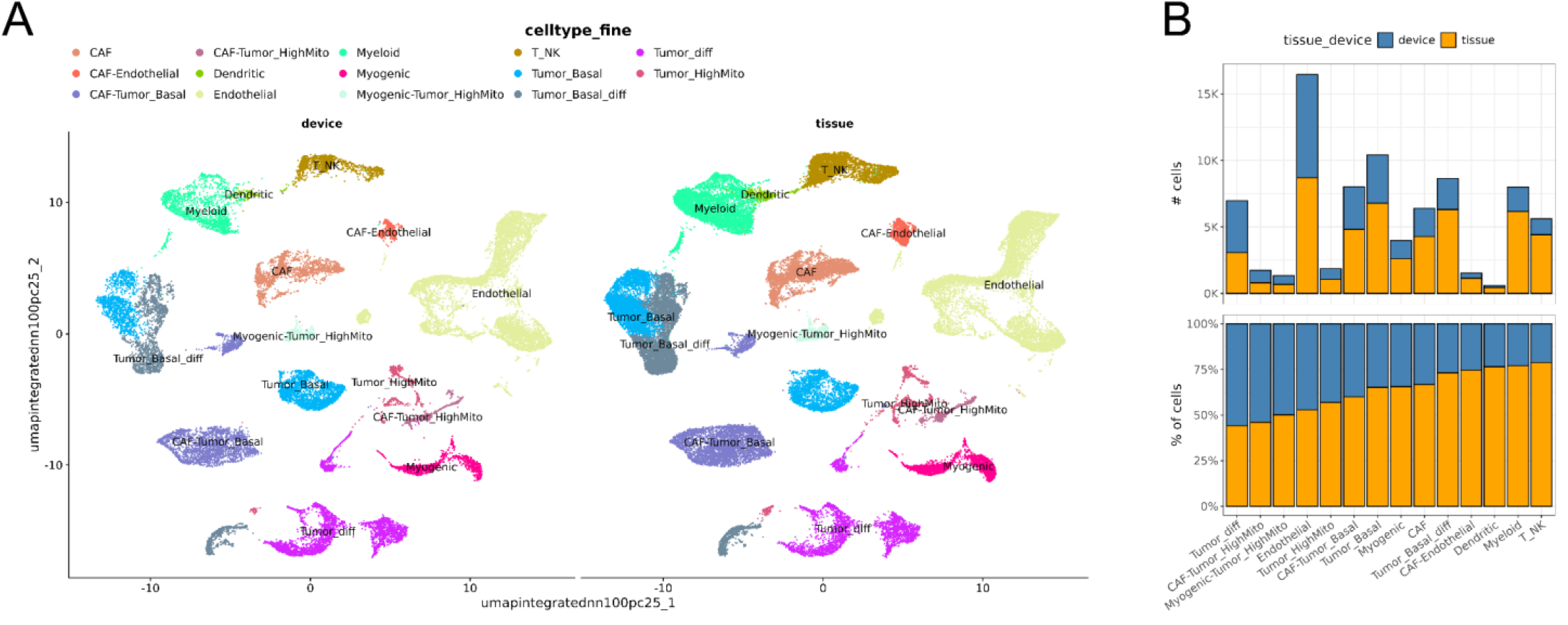
(A) scRNA-seq UMAP representation of cells from MPS (left) and cells from the matched patient tissues (right) for four different patients after data integration using Harmony and STACAS. All profiled samples displayed a high degree of agreement between the patient tissue and MPS made from the same tissue, highlighting the capability of the MPS to maintain transcriptional concordance with the original tissue. (B) Relative abundance plots showing the number and percentage of cells in the MPS (blue) and the tissue (yellow). Each bar in the relative abundance plot corresponds to a cluster in the UMAP.

Cell types were annotated using canonical marker genes listed in **Table 1** and dot plots to verify expression across clusters are shown in **Figure S3**. Preliminary analyses identified subtypes of stromal cells such as fibroblasts, epithelial cells and endothelial cells which were conserved across both tissue and MPS. Additionally, we observed clusters of T and NK cells, along with a smaller population of myeloid and dendritic cells in the MPS (Figure 3B). UCell scores confirm each cell type’s ‘s expression profile **(Figure S4)** Notably, the MPS supported the maintenance of both stromal and immune cell compartments, highlighting the ability of the MPS to maintain a heterogeneous microenvironment reflective of the native tissue composition. The presence of immune populations within the MPS is noteworthy, as the retention of TILs has been a persistent challenge in *in vitro* culture systems and is critical for modeling immune-stromal crosstalk and therapeutic response.

We next investigated the fractional composition of cells in the MPS by plotting the relative abundance of the different clusters across the tissue and MPS **(Figure 3B)**. As expected, cell numbers were higher in the tissue as compared to the MPS, however, the MPS had a higher representation of stromal cells (fibroblasts, and endothelial cells) (>50 %) as compared to the tissue and a lower representation of immune cells (< 30%) relative to the tissue.

These data strongly support the robustness of our digest and cell expansion protocol and establish a strong molecular foundation supporting the use of this MPS for functional patient studies.

### Patient-specific tumor cells elicit differential responses from the endothelial lumens

HNC tumors are known to be highly heterogeneous in their functional behavior across patients, resulting in varied responses to treatment. Based on data from **Figure 3**, we further investigated whether the molecular similarities of the TME in the MPS to the host tissue will translate to functional, phenotypic differences in the MPS such as cell morphology, spheroid invasiveness and endothelial sprouting.

MPS were made from patient cells obtained from two different patients and cultured for 5 days without any treatment. After 5 days, the MPS was stained for viability using Calcein-AM and Propidium Iodide, followed by a Hoechst counter stain for nuclei. As seen in **Figure 4**, MPS from both patients displayed visual differences. Patient #0322 showed confluent, dense endothelial lumens with minimal gaps. The tumor spheroid was largely intact and circular, suggesting a collective cell migration response, and CAF appeared to be rounded and smaller. In contrast, MPS from Patient #1123 displayed angiogenic sprouting in both lymphatic and blood endothelial lumens, with a disperse tumor spheroid indicating a single cell invasive phenotype, coupled with elongated CAF.

**FIgure 4.**
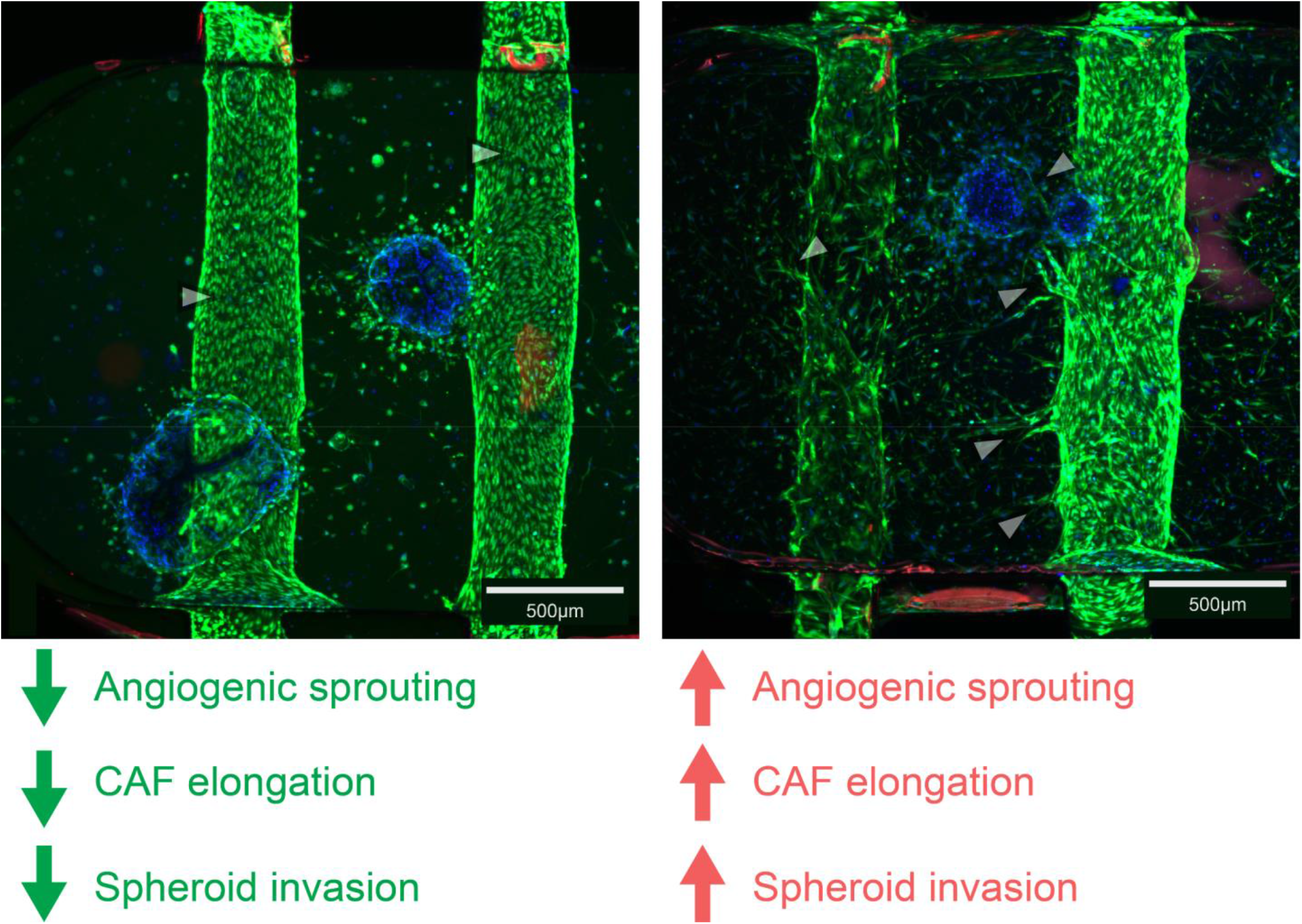
Confocal micrographs of MPS developed from two different matched patient cells. MPS from patient 0322 shows confluent, packed luminal structures with spheroids displaying a collective front type migration profile,. In addition, minimal CAF elongation is observed in this sample. In contrast, cells from patient 1123 induced noticeable endothelial sprouting in both lymphatic and vascular lumens. CAFs appear to be elongated and the tumor cells display a single cell migration behavior.

It is noteworthy that in both MPS, HUVECs and HLECs were used from the same batch of pooled cells from healthy donors. Therefore, any difference in angiogenic sprouting is a function of altered biomechanical signaling from the stromal cells of the patients evaluated in this experiment. Thus, these images show that the MPS captures phenotypic differences between different patients’ tumor cells and these tumor cells continue to uniquely influence the TME.

### HNC MPS captures differential responses of the tumor microenvironment to standard of care chemoradiation

Patients with HNC are most commonly administered radiation therapy and may also receive concurrent platinum-based chemotherapy such as cisplatin depending on the patient risk group. Predicting a patient’s response to therapy remains challenging owing to extensive heterogeneity across patient populations without defined biomarkers. Therefore, to determine whether the MPS can be used to evaluate the effects of chemoradiation, we quantified MPS response to chemoradiation using the SCC1 cell line (a HNC cell line) as the epithelial population, along with primary fibroblasts from a single donor, and pooled HUVECs and HLECs.

The cells were loaded into the MPS and cultured for 24 hours prior to treatment. Cells were treated with Cisplatin diluted in cell culture media, at final concentrations ranging from 7.5 µg mL^−1^ to 2.5 µg mL^−1^. These concentrations were selected based on the plasma trough concentration of cisplatin observed in HNC patients post infusion^39–41^ and cross-validated in 2D cultures **(Figure S5)**. The MPS received a single dose of radiation of 4 Gy in the radiated condition.

After the experiment, cells were stained for viability and imaged using confocal microscopy to evaluate the cellular viability of the epithelial spheroids, fibroblasts, and endothelial lumens using an automated image processing pipeline implemented in Python. As shown in **Figure 5A**, the confocal micrograph was segmented into lumens, spheroids, and tumor stroma following which individual nuclei were identified using StarDist^34,35^. Tumor cells were identified by a DiD membrane dye and nuclei data was used to determine cell viability as fraction of cells expressing propidium iodide, nuclei size, and cell counts per spheroid. In addition, the spheroid area was also recorded. To ensure that any observed differences in our experiments were a result of patient-specific behavior and not experiment-experiment variability, we maintained a control condition with SCC-1 cell line as the malignant cells coupled with selected CAF population. This control condition was evaluated for consistency in spheroid size and viability across patient-based experiments (**Figure S6)** to verify experimental fidelity.

**Figure 5.**
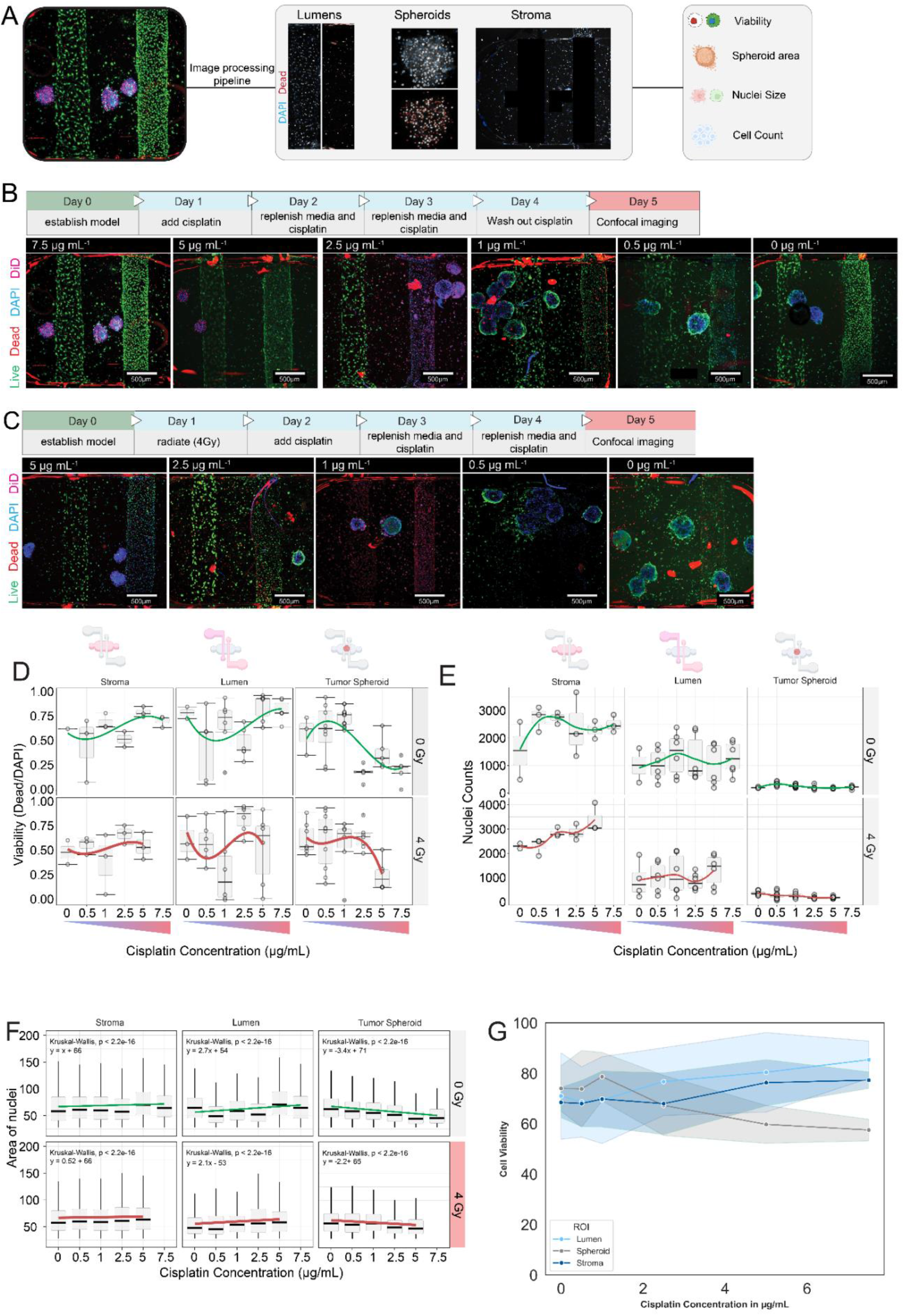
(A) Schematic representation of the image processing steps to extract data from MPS. Images were automatically segmented into stroma and spheroids, and lumen ROIs were manually defined. An automated script extracted spheroid areas, cell counts, nuclei area, and viability. (B, C) Treatment timelines for evaluating cisplatin dose-responses without and with radiation in the MPS. Corresponding panels under each timeline show representative confocal micrographs of the TME in the MPS for each treatment condition. (D) Boxplots showing cell viability by region in the MPS. Local regression trendline with standard error is shown for each region condition. (E) Boxplots showing nuclei counts per spheroid for each condition with local regression trendlines. (F) Boxplots showing the area for individual nuclei in different regions of the MPS with linear trendlines. (G) Cell viability percentages were aggregated for both radiated and unirradiated conditions to visualize the effect of cisplatin across regions.

**Figures 5B and 5C** show the timeline for the dose response experiments without and with radiation respectively along with representative confocal micrographs which have been max projected for visualization. Each micrograph shows both the blood endothelial lumen and lymphatic lumens along with tumor spheroids, and stroma cell populations.

**Figure 5D** shows the cell viability across the different compartments of the device. Each point represents an individual spheroid, lumen, or stromal compartment. As expected, the tumor spheroids displayed a sigmoidal response to cisplatin concentration with a drop in viability at 2.5 µg mL^−1.^ In contrast, the endothelial lumens and stromal cells showed a dip in viability that correlated with increased viability of tumor spheroids, suggesting a negative influence of the tumor spheroid on the microenvironment. In the non-radiated condition, we can see that the lumen viability in the control condition is ~76%. When cisplatin concentration is increased from 0 to 1 µg mL^−1^, the endothelial viability decreases to ~40% before increasing to baseline again at higher cisplatin concentrations. A similar trend is observed in the stromal compartment - where the fibroblast viability recovers at the same cisplatin concentration at which the tumor viability decreases. We posit that at intermediate cisplatin concentrations (0.5, 1 µg mL^−1^), endothelial cells and fibroblasts experience toxicity from the cisplatin in addition to pro-inflammatory cytokines and competition for nutrients from tumor cells. However, when the model is exposed to higher concentrations of cisplatin, epithelial viability reduces disproportionately as the epithelial cells have a higher proliferative capacity and are more likely to respond to DNA damage from cisplatin. Consequently, the rapid reduction of the epithelial fraction reduces competition for nutrients, and since the endothelial cells and fibroblasts are slower proliferating cells, they exhibit lower sensitivity to DNA damage, resulting in their recovery.

Nuclei count across compartments provide a measure of proliferative capacity in response to treatment **(Figure 5E)**. In the non-radiated condition, nuclei count in the tumor spheroid, lumen, and stroma remained consistent across increasing cisplatin doses. However, upon the addition of radiation, nuclei count in the tumor spheroids decreased with increasing cisplatin concentration, while stromal and lumen nuclei count increased. This inverse relationship may reflect competitive release within the confined MPS microenvironment: as tumor cells are eliminated by chemoradiation, reduced competition for nutrients and space may permit expansion of non-malignant populations.

Orthogonally, nuclei size is often used as a marker of cell health with smaller nuclei associated with apoptotic cells. In the tumor spheroid, nuclei area was found to decrease with increasing cisplatin dose (slope = −3.4, −2.2). However, the slope nuclei area of endothelial cells displayed a positive slope in response to cisplatin dose (2.7, 2.1). Stromal cells displayed near constant nuclei size in response to treatment.

Together, these data demonstrate that the MPS enables spatially resolved quantification of treatment response across distinct cellular compartments within the tumor microenvironment. By coupling the MPS with an automated image processing pipeline, we extracted four independent functional readouts comprising viability, nuclei count, nuclei size, and spheroid area from each compartment within a single device. This approach reveals compartment-specific dynamics, such as the reciprocal relationship between tumor and stromal viability, that would not be accessible in conventional monoculture systems. The ability to resolve these interactions at the compartment level positions the MPS as a platform for dissecting how individual populations within the TME respond to treatment.

### HNC MPS generates a hypoxic cell culture environment which reduces radiation response

Hypoxia is a hallmark of head and neck cancer (HNC) tumors and a principal driver of radioresistance, as the reduced availability of oxygen attenuates the fixation of radiation-induced DNA damage by reactive oxygen species (ROS)^25–27^. We assessed whether the MPS recapitulates the hypoxic conditions characteristic of the HNC tumor microenvironment (TME) and whether this influences radiation response relative to conventional two-dimensional (2D) monolayer cultures.

To quantify the degree of hypoxia within MPS-cultured tumor spheroids, we employed a fluorescent hypoxia indicator dye and compared signal intensity between spheroids maintained in the MPS and 2D monolayer cultures following 24-hour incubation under hypoxic (1% O_2_) and normoxic conditions. Spheroids cultured within the MPS exhibited pronounced hypoxia even under normoxic culture conditions, with a 12-fold increase in mean fluorescence intensity relative to 2D baseline cultures **(Figure 6A, B)**. This intrinsic hypoxia likely reflects oxygen diffusion limitations within the three-dimensional architecture of the spheroid and the surrounding MPS microenvironment, consistent with the oxygen gradients observed in solid tumors *in vivo*^42^.

**Figure 6.**
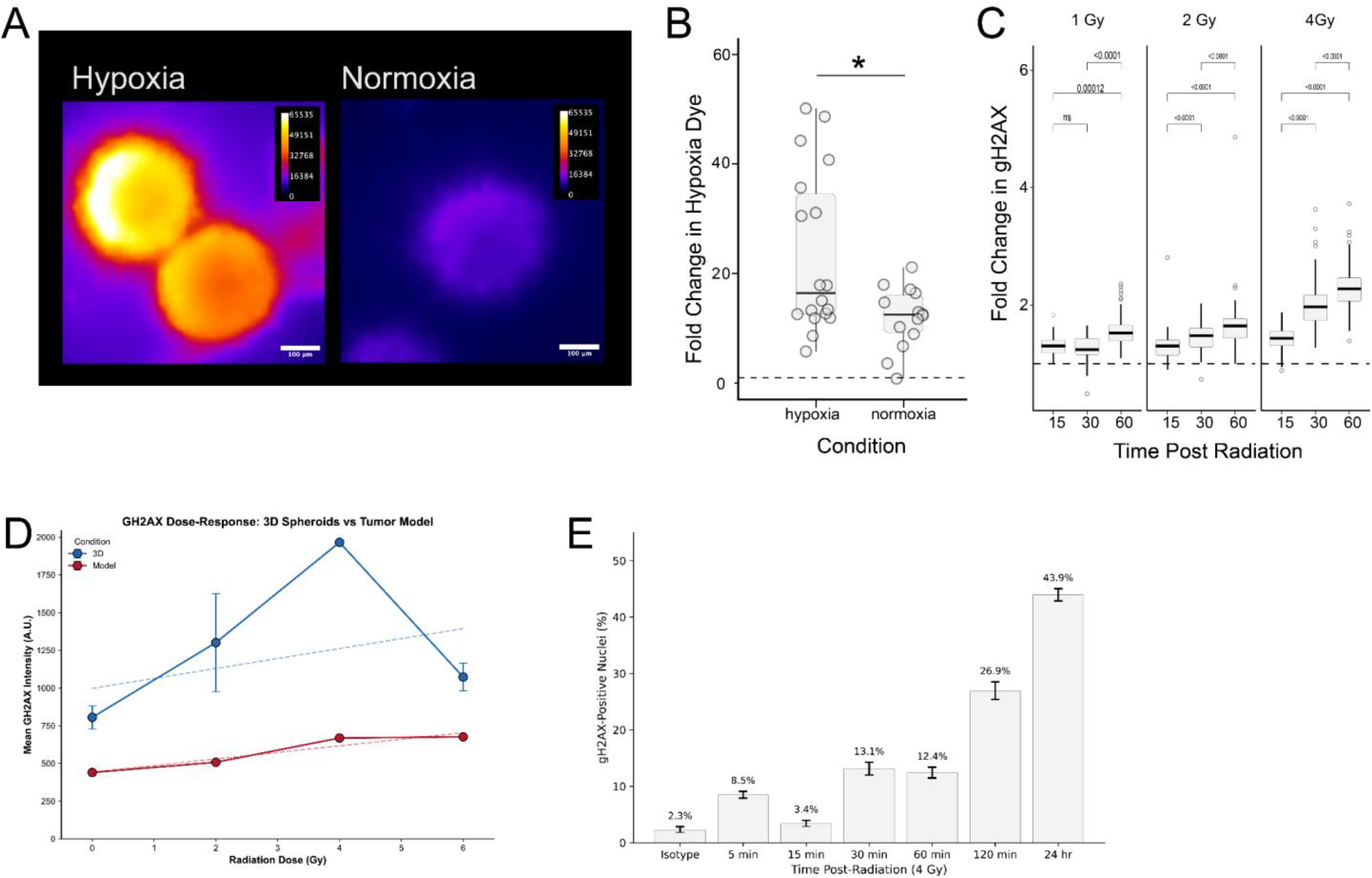
(A) Intensity of hypoxia marker within spheroids incubated inside a hypoxic chamber as compared to a normoxic chamber. (B) Fold change in hypoxia dye intensity in both normoxic and hypoxic cultured spheroids compared to a 2D plate culture. (C) Fold change in gH2AX expression over time in minutes after radiation in 2D at radiation doses of 1 Gy, 2 Gy, and 4 Gy. A consistent increase in Gh2AX expression can be observed in 2D. (D) GH2AX expression in 3D (blue) and MPS (red) with increasing radiation dose shows MPS to be less sensitive to increasing dose, reinforcing the role of hypoxia and stromal cells in modulating radiation response. (E) Time course of gH2AX expression in MPS after radiation with 4Gy.

We next investigated whether this hypoxic milieu conferred differential radiation sensitivity. 2D monolayer cultures of SCC1 cells were irradiated at doses of 1 Gy, 2 Gy, and 4 Gy, and cells were fixed at 15, 30, and 60 minutes post-exposure and immunostained for γH2AX, a well-established marker of DNA double-strand breaks (DSBs)^43,44^. Across all dose levels, γH2AX signal increased over the 60-minute observation window, consistent with ongoing DSB induction and the recruitment of DNA damage response machinery within this early time frame **(Figure 6C)**.

To assess the effect of radiation dose on DSB burden across culture formats, we compared γH2AX expression in 3D spheroids and MPS constructs exposed to escalating radiation doses. MPS cultures exhibited attenuated γH2AX induction relative to spheroids cultured outside the MPS at equivalent doses **(Figure 6D)**, suggesting that the MPS microenvironment encompassing both the hypoxic niche and the presence of stromal cellular constituents confers a radioprotective effect that reduces the accumulation of unresolved DSBs ^45,46^.

To further dissect DSB repair kinetics within the MPS, SCC1-based constructs were irradiated at 4 Gy and γH2AX expression was monitored over a 24-hour time course. A rapid increase in γH2AX signal was observed within 5 minutes post-irradiation, followed by a sustained, gradual rise in expression extending to 24 hours **(Figure 6E)**. This biphasic response is consistent with the well-described two-component model of DSB repair, in which fast repair, which is predominantly mediated by non-homologous end joining (NHEJ), resolves the majority of lesions within 15-20 minutes, while a slow repair component, likely involving homologous recombination or repair of complex, clustered lesions, proceeds over several hours. We attribute the protracted repair kinetics observed in the MPS relative to 2D cultures to the influence of the three-dimensional hypoxic microenvironment on DNA damage resolution.

Taken together, these findings demonstrate that the MPS establishes a constitutively hypoxic microenvironment that modulates radiation-induced DNA damage and repair dynamics in a manner reflective of the native HNC TME, thereby providing a more physiologically relevant platform for evaluating radiation response than conventional 2D culture systems.

### Patient-specific MPS recapitulate differential treatment sensitivity and identify potential predictors of clinical outcome

To evaluate the capacity of the MPS platform to capture patient-specific treatment responses, we constructed MPS from tumor specimens obtained from multiple HNC patients and subjected them to treatment regimens comprising fractionated radiation (0-32 Gy, delivered in two fractions) alone or in combination with cisplatin. The dose of radiation was selected to simulate an equivalent dose in 2 Gy (EQD_2_) of 70 Gy - the standard dose of radiation delivered to patients over a span of 30 days - with 32 Gy delivered over two fractions equivalent to an EQD_2_ of 69.33 Gy, assuming α/β = 10 for HNC^47^.

Three orthogonal readouts, spheroid area, cell viability, and cell count, were quantified as a function of treatment intensity for each patient **(Figure. 7A–C)**. Across all patients, dose-dependent reductions in spheroid area, viability, and cell count were observed with escalating radiation dose, and combination with cisplatin further augmented treatment effect, consistent with the established role of cisplatin as a radiosensitizer in HNC. Importantly, the magnitude and kinetics of these responses varied across patients, reflecting the interpatient heterogeneity in treatment sensitivity that is well documented clinically but poorly captured by conventional in vitro models.

**Figure 7.**
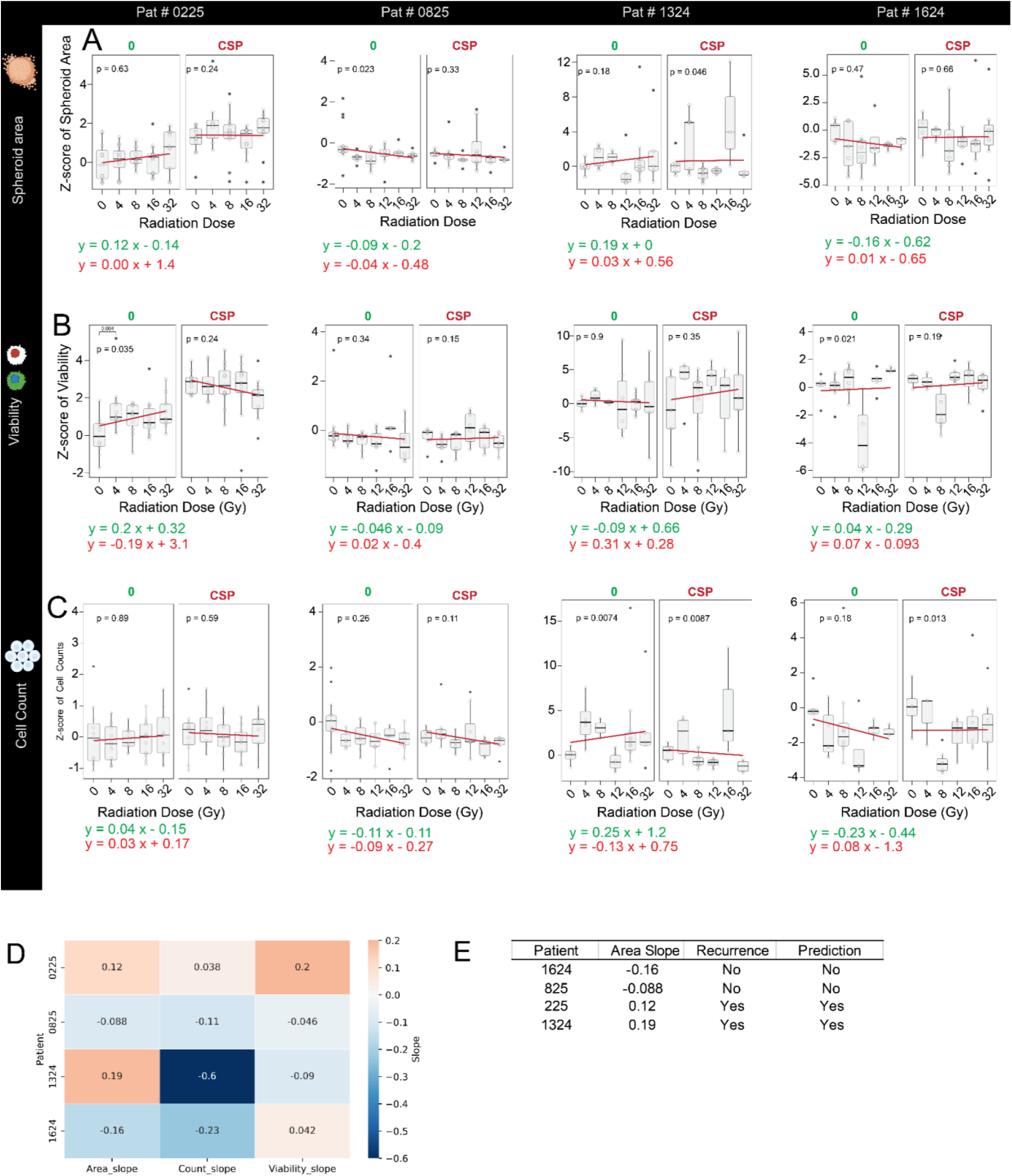
Plots showing treatment outcomes from patient specific MPS treated with radiation (0-32Gy, 2 fractions, ± Cisplatin). Each column represents a single patient as indicated in the header. (A) Spheroid as a function of treatment. (B) Cell viability as a function of treatment. (C) Cell count as a function of treatment. (D) Heatmap with slopes of linear regressions fit on each readout for each patient. (E) The area slope appears to be a strong predictor of patient outcome (where outcome is defined as recurrence at a follow-up visit) with a negative slope suggesting that the patient did not recur and a positive slope indicating recurrence.

To distill these multiparametric response profiles into a quantitative framework, we fit linear regressions to each readout as a function of treatment intensity for each patient and extracted the corresponding slopes as summary metrics of treatment sensitivity **(Figure. 7D)**. Hierarchical visualization of these slopes across patients and readouts revealed patient-specific response signatures, with some patients exhibiting uniformly steep negative slopes which were indicative of robust treatment sensitivity and others displaying attenuated or positive slopes suggestive of relative treatment resistance.

Among the three readouts evaluated, the slope of the spheroid area as a function of treatment emerged as a candidate predictor of clinical outcome. In this preliminary cohort, patients whose MPS demonstrated a negative area slope and thus reflected dose-dependent tumor shrinkage did not experience disease recurrence at clinical follow-up (>5 months post treatment), whereas patients with a positive area slope suggesting maintained or increasing spheroid area despite treatment did subsequently recur **(Figure. 7E)**. While these observations are derived from a limited patient cohort and are therefore hypothesis-generating rather than definitive, the concordance between ex vivo MPS response and clinical outcome is noteworthy and suggests that the MPS platform may capture biologically meaningful aspects of individual treatment sensitivity.

These proof-of-concept data provide an initial indication that patient-derived MPS, when coupled with quantitative response metrics, may hold potential as a functional precision oncology tool for predicting treatment outcomes in HNC. Ongoing studies incorporating expanded patient cohorts and the development of robust statistical models will be essential to validate these preliminary findings and to establish the sensitivity, specificity, and clinical utility of MPS-derived response signatures as predictive biomarkers.

## Discussion

The capacity to predict individual patient response to therapy remains a central unmet need in HNC management. In this study, we developed a patient-specific MPS that recapitulates critical features of the HNC TME and demonstrated its potential for evaluating individual treatment response to chemoradiation.

Our MPS incorporates diverse cell populations which include patient-derived tumor epithelial cells, cancer-associated fibroblasts, multiple tumor-infiltrating leukocyte subtypes, blood endothelial cells, and lymphatic endothelial cells, within a 3D collagen-fibronectin hydrogel containing perfusable vascular and lymphatic lumens. In comparison, existing organoid and spheroid-based platforms typically lack the stromal and vascular compartments that are critical mediators of treatment response. The inclusion of lymphatic endothelium is significant, as lymphatic vessels are involved in HNC metastasis and immune trafficking yet are largely absent from existing in vitro models. Our prior work has demonstrated that the HNC TME is sensitive to lymphatic vessel presence^19^, which can promote tumor cell migration, motivating the inclusion of lymphatic endothelial cells in the MPS.

The fidelity of an *in vitro* model hinges on its capacity to preserve the molecular identity of the cells it seeks to represent. scRNA-seq analysis across patients demonstrated transcriptomic concordance between MPS-cultured cells and matched primary tumor tissue, with 10 transcriptionally distinct cell populations from both sources co-localizing in shared UMAP embedding space. The preservation of TILs is significant given the increasing clinical relevance of the immune compartment in HNC, both as a prognostic indicator and as a determinant of response to immunotherapy^48–50^. The concurrent maintenance of epithelial, stromal, endothelial, lymphatic, and immune populations within a single MPS construct, with demonstrated transcriptomic concordance to the source tissue, has not been previously reported for HNC. While we limited our study to chemoradiation because all the patients in our cohort only underwent radiation with or without chemotherapy, recently neoadjuvant immunotherapy (pembrolizumab) has been approved as a new standard of care option^51^. Given the inclusion of immune cells and lymphatics in the MPS, this model also holds strong potential to report immunotherapy responses in the future.

The endothelial and lymphatic cells used in our MPS were derived from pooled healthy donors rather than from the patient tissue, owing to the limited size of surgical specimens and the inability to isolate sufficient autologous endothelial populations. Despite this limitation, we observed patient-specific differences in endothelial behavior, including differential angiogenic sprouting and lumen morphology, driven entirely by the patient-derived stromal compartment. These findings suggest that tumor-stromal paracrine signaling is a dominant driver of endothelial phenotype within the MPS, partially mitigating the limitation of using allogeneic endothelial cells and reinforcing the importance of the stromal compartment in governing TME behavior. A patient-specific response to endothelial sprouting has also been observed by us in previous studies^10,23,52^. Synergistically, our media optimization studies also highlight that co-culture itself was a stronger driver of cell viability than media composition alone, highlighting the importance of paracrine signaling within the TME and providing further justification for the use of multicellular MPS over simpler culture formats.

The capacity of the MPS to recapitulate a hypoxic microenvironment characteristic of HNC tumors is a notable finding. Hypoxia altered radiation response with attenuated γH2AX induction relative to 3D spheroids cultured outside the MPS, and the kinetics of DNA repair within the MPS followed a biphasic profile, with a rapid initial component followed by slow repair extending to 24 hours, more consistent with in vivo radiobiology than the repair dynamics observed in 2D monolayers^45,46^. The ability to generate hypoxia without specialized chambers simplifies functional testing workflows. However, the relative contributions of hypoxia, stromal cell-mediated radioprotection, and ECM-mediated signaling to the radiation response remain to be formally deconvolved.

The cell-compartment specific analysis revealed an interaction between tumor and stromal populations in response to chemoradiation. Endothelial and fibroblast viability initially decreased at intermediate cisplatin concentrations before recovering at higher doses, coincident with the collapse of tumor cell viability, suggests competitive dynamics within the microenvironment that are not captured by monoculture systems. This reciprocal relationship between malignant and non-malignant compartments has implications for understanding treatment-induced remodeling of the TME.

The translational potential of the MPS was evaluated through patient-specific models treated with fractionated chemoradiation regimens. Across the cohort, patients exhibited heterogeneous dose-response profiles as assessed by spheroid area, cell viability, and cell count, three orthogonal readouts that capture complementary aspects of treatment response. The extraction of linear regression slopes from these dose-response curves provided a quantitative summary metric for each patient, and among the readouts evaluated, the slope of the spheroid area emerged as a candidate predictor of clinical outcome. Patients with negative area slopes, reflecting dose-dependent tumor shrinkage, did not experience disease recurrence at follow-up up to 12 months, while those with positive slopes subsequently recurred within 12 months. The potential superiority of area as a predictive readout over viability or cell count may reflect the fact that spheroid area integrates multiple biological processes (proliferation, death, compaction, and invasion) into a single morphometric measurement, whereas viability and cell count capture more isolated aspects of the treatment response.

We emphasize that these observations, while novel, are preliminary and hypothesis-generating. The limited patient cohort precludes formal statistical validation, and additional confounding variables (including differences in tumor staging, anatomical subsite, and treatment history) have not been systematically controlled for. Establishing the predictive utility of MPS-derived response metrics will require further studies with expanded cohorts, blinded analysis, and statistical modeling to determine sensitivity, specificity, and positive and negative predictive values. Furthermore, the current experimental timeline spans approximately two to three weeks, which may limit clinical applicability in settings where treatment decisions must be made rapidly. Ongoing efforts are focused on streamlining the workflow to reduce turnaround time and on transitioning to higher-throughput device formats^53^.

Despite these limitations, the data presented here establish, for the first time, proof-of-concept for a patient-specific HNC MPS that integrates the cellular, structural, and microenvironmental complexity of the native TME and generates quantitative, patient-specific functional readouts in response to clinically relevant treatment regimens. The concordance between MPS response and clinical outcomes, while preliminary, provides rationale for continued development and validation of this platform as a functional precision oncology tool. Looking forward, the integration of additional readouts, including secretomic profiling, spatial transcriptomics, and real-time metabolic monitoring, may further enhance the predictive resolution of the system. Moreover, modular modifications to the LumeNEXT platform and possibility of higher throughput studies positions it for immunotherapy and targeted therapy testing, areas of growing clinical importance in HNC that are currently underserved by existing functional assay platforms.

In summary, these data support the potential of MPS-based functional assays as a complementary approach to molecular profiling in precision oncology for head and neck cancer.

## Acknowledgements

This project was supported by the Specialized Program of Research Excellence (SPORE) program, through the NIH National Institute for Dental and Craniofacial Research (NIDCR) and National Cancer Institute (NCI), grant P50CA278595, and Carbone Cancer Center Support Grant NIH P30CA014520. The content is solely the responsibility of the authors and does not necessarily represent the official views of the NIH. We also acknowledge the use of resources afforded by the University of Wisconsin Optical Imaging Core, Small Animal Imaging and Radiotherapy Facility, Small Molecule Screening Facility, and the University of Wisconsin Biotechnology Center. We thank the Wisconsin Head and Neck SPORE Pathology and Biospecimen core for access to patient tissue.

