## Supplementary Data for "A Head and Neck Cancer Patient-Specific Microphysiological System for Predicting Response to Chemoradiation"

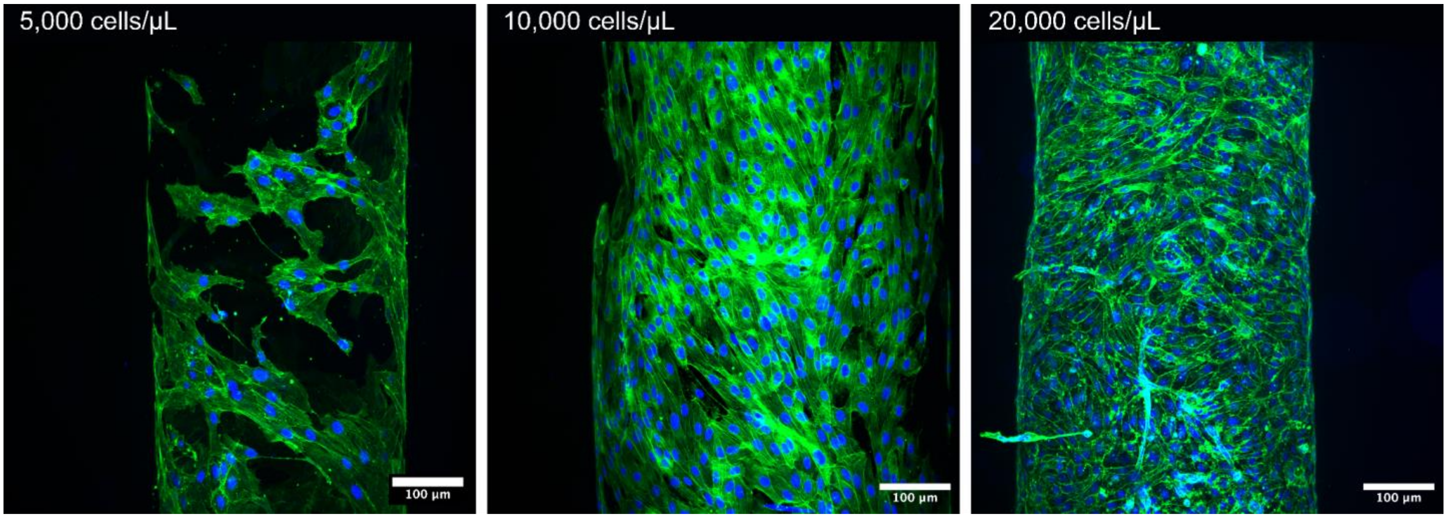


**Figure S1:** Image showing max projections of lumens seeded at cell densities ranging of 5,000 - 20,000 cells $\mu L^{-1}$.

| Media type | Pneumacult | EBM-EGM2 | FM |
|---|---|---|---|
| 1 | 0 | 100 | - |
| 2 | 20 | 80 | - |
| 3 | 40 | 60 | - |
| 4 | 60 | 40 | - |
| 5 | 80 | 20 | - |
| 6 | 100 | 0 | - |
| 7 | 0 | 0 | 100 |
| 8 | 10 | 10 | 80 |
| 9 | 20 | 20 | 60 |
| 10 | 30 | 30 | 40 |
| 11 | 40 | 40 | 20 |
| 12 | 50 | 50 | 0 |

**Figure S2:** Table showing the media formulations as a percentage of each component for different media tested. In each case, equal amount of basal media were used, with different ratios of added supplements.

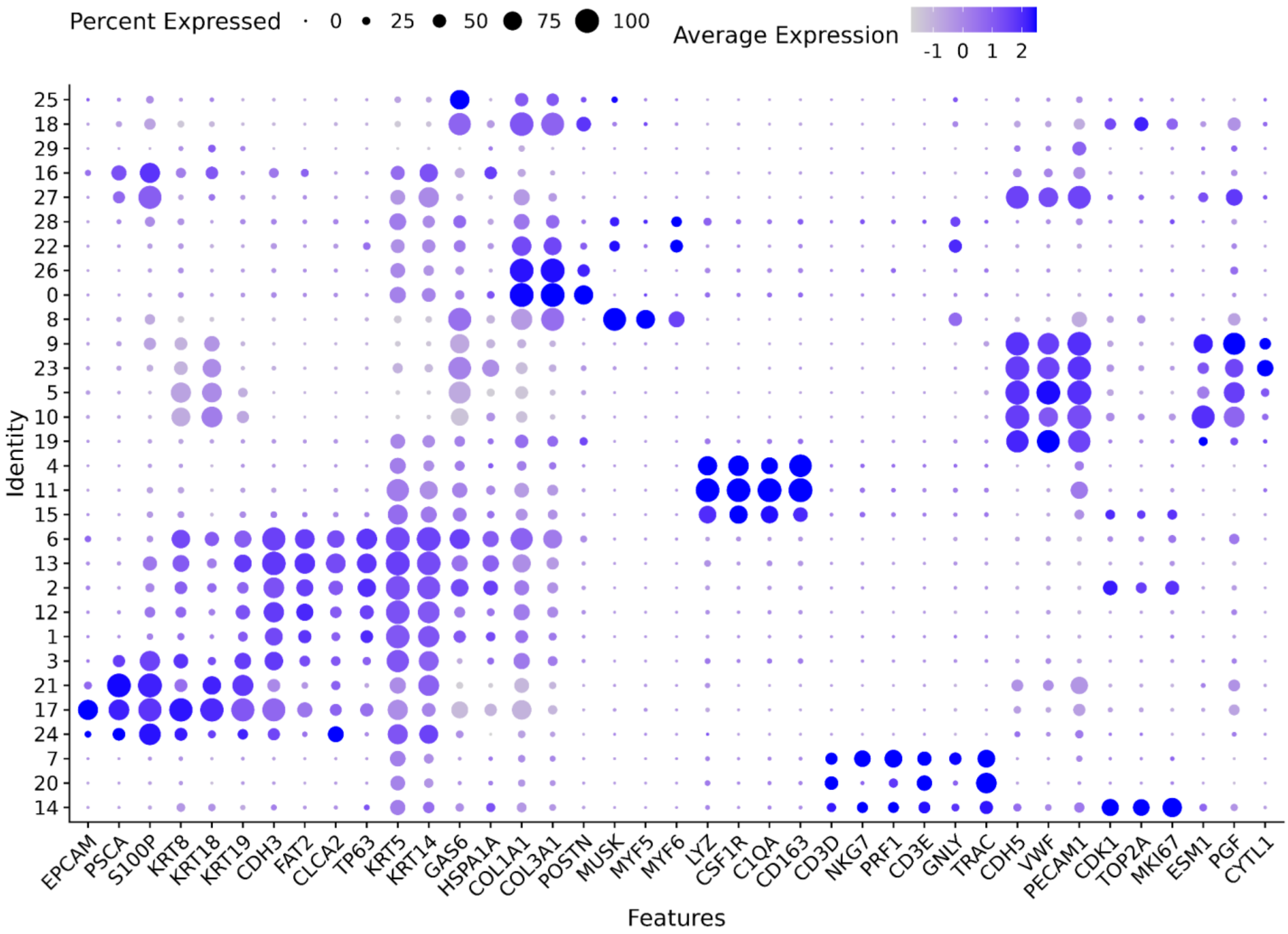

Percent Expressed
0
25
50
75
100
Average Expression
-1 0 1 2
Identity
25
18
29
16
27
28
22
26
0
8
9
23
5
10
19
4
11
15
6
13
2
12
1
3
21
17
24
7
20
14
EPCAM PSCA S100P KRT8 KRT18 KRT19 CDH3 FAT2 CLCA2 TP63 KRT5 KRT14 GAS6 HSPA1A COL1A1 COL3A1 POSTN MUSK MYF5 MYF6 LYZ CSF1R C1QA CD163 CD3D NKG7 PRF1 CD3E GNLY TRAC CDH5 VWF PECAM1 CDK1 TOP2A MKI67 ESM1 PGF CYTL1
Features

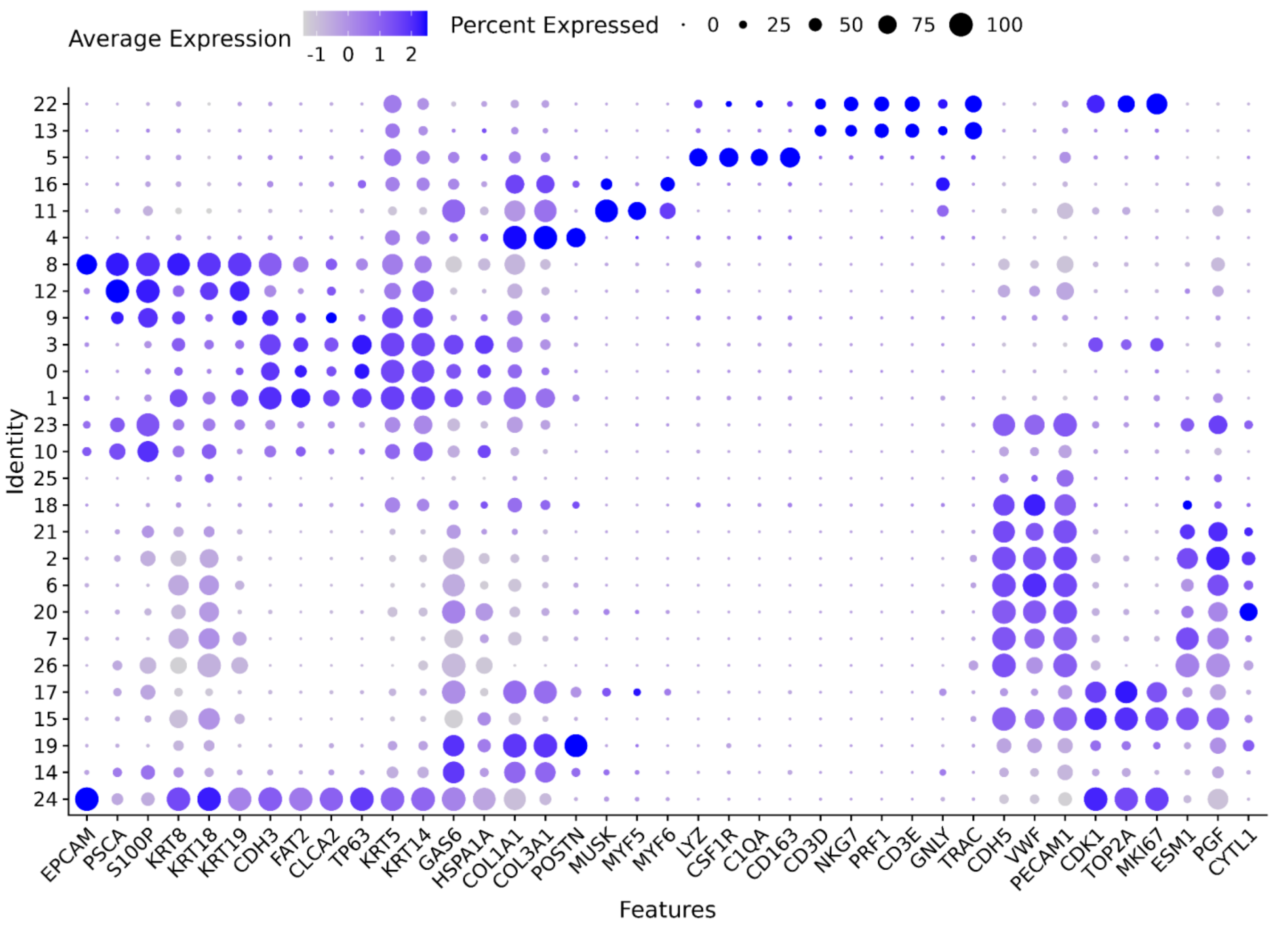


**Figure S3:** Dot plots showing cell type annotation for the integrated sc RNA-seq tissue (top) and MPS devices (bottom) datasets. The columns correspond to the gene markers in Table 1 and the rows are the dataset specific clusters. The dots show the avg. expression of each gene per cluster and the percentage of cells in the clusters showing gene expression.

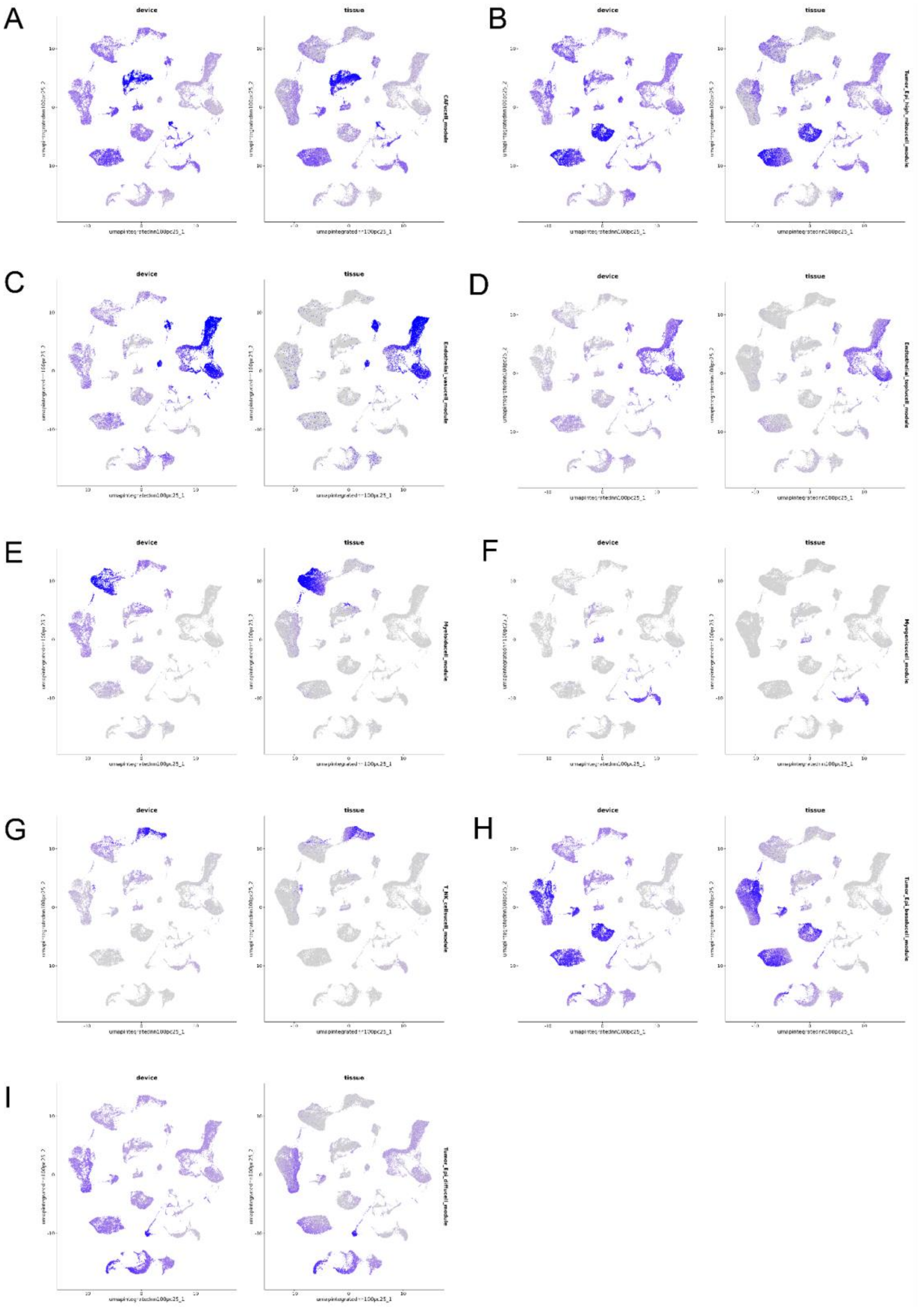


**Figure S4:** UMAP representations coloured by celltype module indexes computed with Ucell: (A) CAF, (B) Tumor_epi_high_mito, (C) Endothelial_vasc, (D) Endothelial_tipI, (E) Myeloid_cells, (F) Myogenic_cells, (G) T_NK_cells, (H) Tumor_epi_basal_cells, (I) Tumor_epi_diff_cells, and (J) Dendritic cells.

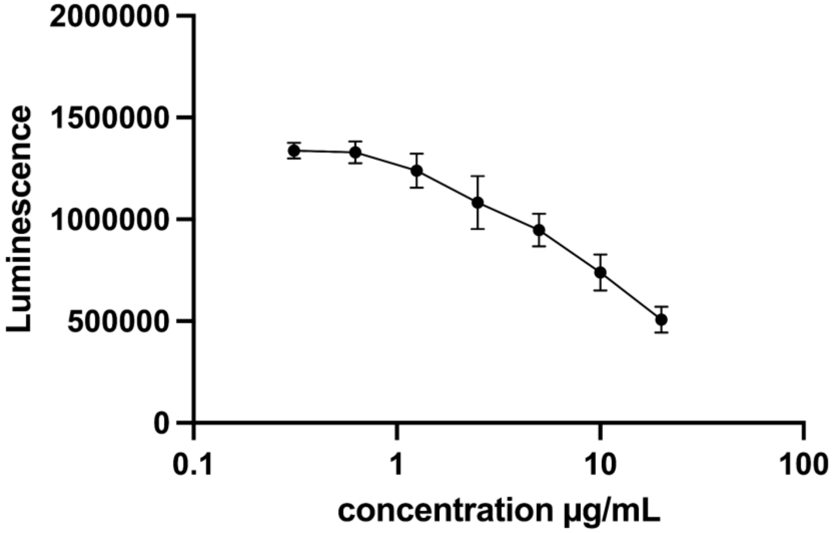


**Figure S5:** Response of HNC epithelial cells to cisplatin in 2D.

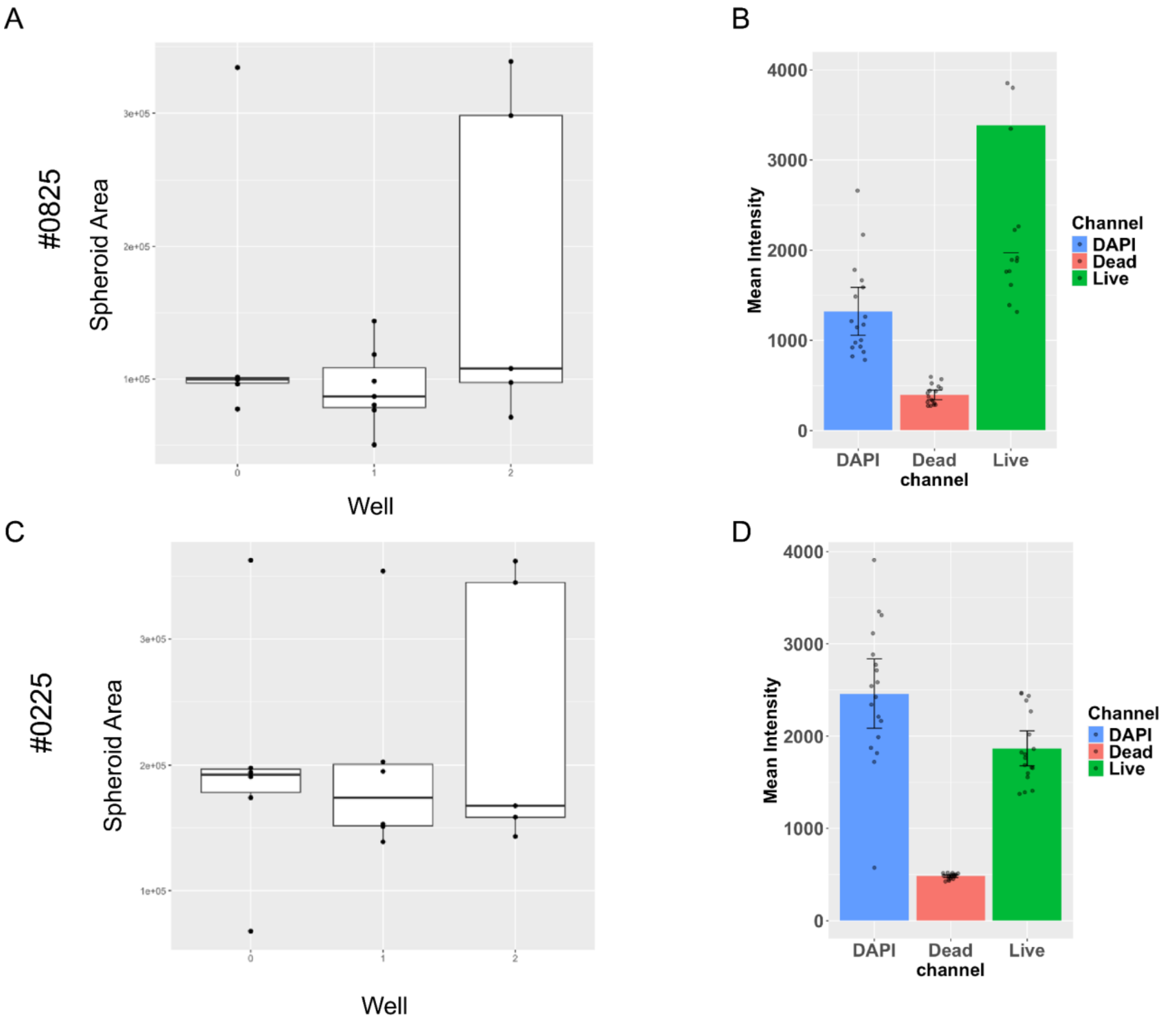


**Figure S6:** Plots showing representative example measurements of spheroid area and staining intensity values for SCC1 control spheroids which were cultured simultaneously alongside patient-specific devices to test for run-run variability across experiments. (A) Spheroid area for SCC-1 spheroids cultured alongside #0825 samples, (B) Staining intensity values (DAPI, propidium iodide and calcein-AM) for SCC-1 spheroids cultured alongside #0825 samples. (C-D) Spheroid areas and staining intensities for SCC-1 spheroids cultured alongside #0225 samples.
